# Widely used commercial ELISA does not detect preHP-2, but recognizes properdin as a potential second member of the zonulin family

**DOI:** 10.1101/157578

**Authors:** Lucas Scheffler, Alyce Crane, Henrike Heyne, Anke Tönjes, Dorit Schleinitz, Christian H. Ihling, Michael Stumvoll, Rachel Freire, Maria Fiorentino, Alessio Fasano, Peter Kovacs, John T. Heiker

## Abstract

**BACKGROUND:** There is increasing evidence for the role of impaired intestinal permeability in obesity and associated metabolic diseases. Zonulin is an established serum marker for intestinal permeability and identical to pre-haptoglobin2. Here, we aimed to investigate the relationship between circulating zonulin and metabolic traits related to obesity.

**METHODS:** Serum zonulin was measured by using a widely used commercial ELISA kit in 376 subjects from the metabolically well-characterized cohort of Sorbs from Germany. In addition, haptoglobin genotype was determined in DNA samples from all study subjects.

**RESULTS:** As zonulin concentrations did not correlate to the haptoglobin genotypes, we investigated the specificity of the zonulin ELISA assay using antibody capture experiments, mass spectrometry and Western blot analysis. Using serum samples that gave the highest or lowest ELISA signals, we detected several proteins that are likely to be captured by the antibody in the present kit. However, none of these proteins corresponds to pre-haptoglobin2. We used increasing concentrations of recombinant pre-haptoglobin 2 and complement C3 as one of the representative captured proteins and the ELISA kit did not detect either. Western blot analysis using both the polyclonal antibodies used in this kit and monoclonal antibodies rose against zonulin showed a similar protein recognition pattern but with different intensity of detection. The protein(s) measured using the ELISA kit was (were) significantly increased in patients with diabetes and obesity and correlated strongly with markers of the lipid and glucose metabolism. Combining mass spectrometry and Western blot analysis using the polyclonal antibodies used in the ELISA kit, we identified properdin as another member of the zonulin family.

**CONCLUSIONS:** Our study suggests that the zonulin ELISA does not recognize pre-haptoglobin 2, rather structural (and possibly functional) analogue proteins belonging to the mannose-associated serine protease family, with properdin being the most likely possible candidate.

## Introduction

The “intestinal barrier” is an established term, defined as a functional entity separating the gut lumen from the inner host, and consisting of mechanical, humoral, immunological, muscular and neurological elements. Intestinal barrier dysfunction is a characteristic feature of pathological states such as inflammatory bowel disease, celiac disease, nonalcoholic steatohepatitis and ulcerative colitis (1, 2). There is also emerging evidence for the role of impaired intestinal permeability in metabolic diseases including obesity and type 2 diabetes (T2D) (3–5). It has been hypothesized that gut bacteria and bacterial endotoxins may disrupt the intestinal barrier resulting in the so called “leaky gut” (4, 6). The leakage of toxins, bacterial components or even live bacteria and their transfer to target organs such as adipose tissue might contribute to the development of obesity and T2D (6, 7). Indeed, numerous studies in mouse models have demonstrated that changes in the gut microbiota can alter the gut permeability and lead to an endotoxemia-induced inflammation in adipose tissue, and ultimately to obesity (3, 8, 9). Results from experimental mouse models are supported by studies in humans by showing an increase in circulating endotoxin levels and circulating bacterial DNA in obese/diabetic patients, likely due to an increased intestinal permeability in affected subjects (10, 11).

Intestinal barrier transport is mainly regulated by structures of the paracellular pathway called tight junctions which form barriers between epithelial cells and regulate the transport of ions and small molecules across the intestinal lumen. Intestinal permeability is a functional feature of the intestinal barrier. It can be measured by analyzing flux rates of inert molecules across the intestinal wall as a whole or across wall components (1). The gold standard for assessment of intestinal permeability in vivo is an assay combining indigestible large and small oligosaccharides, such as lactulose and mannitol; the larger oligosaccharide, lactulose, is only transported via the paracellular pathway, whereas the smaller oligosaccharide, mannitol, is taken up freely over the intestinal barrier via the transcellular route. However, these oligosaccharide assays are expensive, laborious, poorly reproducible, and time consuming. Therefore, identifying appropriate biomarkers for intestinal permeability is highly desirable. Zonulin has been identified as a tight junction regulating protein which is, functionally, the human counterpart of the Vibrio cholera endotoxin zonula occludens toxin (12, 13). Precisely, subsequent studies recognized zonulin as the precursor of haptoglobin 2 (pre-HP2) (14). Haptoglobin is a well-known protein involved in scavenging hemoglobin, whereas the function of its precursor is largely unknown. Haptoglobin is first synthesized into a single chain precursor protein, which is cleaved into a light N-terminal α-chain and heavy C-terminal β-chain. An exon duplication of exons 3 and 4 of the haptoglobin gene differentiates the HP1 from the HP2 allele. Due to this exon duplication the HP2 α-chain is 1.7kb longer than in the HP1 allele. Haptoglobin is active as tetramer consisting of 2 α- and 2 β-chains linked by disulfide bonds, resulting in three possible genotypes: homozygous HP1/1 and HP2/2 as well as heterozygous HP1/2 (15, 16). About 15 % of the Caucasian population is homozygous for HP (16, 17). Zonulin as pre-HP2 reversibly opens tight junctions and is upregulated in diseases such as celiac disease and type 1 diabetes (T1D) (14, 18). Serum zonulin concentrations are also increased in T2D and obesity (19–21) and strong correlations were observed with various metabolic markers, including fasting plasma glucose, IL-6, HDL, and triglyceride (TG) levels (19–21).

Here, we aimed at characterizing the relationship between circulating serum zonulin and traits related to obesity in a metabolically well-characterized cohort of Sorbs from Germany. To measure zonulin, we used the commercially available ELISA kit (Immundiagnostik, Bensheim, Germany). In addition, we determined the haptoglobin genotypes in the entire cohort. Due to a lack of correspondence between the observed circulating zonulin concentrations and the haptoglobin genotypes in our study cohort, we further investigated the possible identity of the product captured by the commercial ELISA assay. We found that the ELISA kit used in the present study does not detect purified pre-HP2 but rather targets one or more proteins from a range of candidate molecules possibly structurally and functionally related to zonulin. Our data also showed that protein concentrations measured by this ELISA correlated with parameters of obesity and related metabolic traits.

## Materials and Methods

### Study subjects

All subjects are part of a sample from an extensively clinically characterized population from Eastern Germany, the Sorbs (22–24). Extensive phenotyping included standardized questionnaires to assess past medical history and family history, collection of anthropometric data (weight, height, waist-to-hip ratio (WHR)), and an oral glucose tolerance test. Glucose was assessed by the Hexokinase method (Automated Analyzer Modular, Roche Diagnostics, Mannheim, Germany) and serum insulin was measured using the AutoDELFIA Insulin assay (PerkinElmer Life and Analytical Sciences, Turku, Finland). Total serum cholesterol and TG concentrations were measured by standard enzymatic methods (CHOD-PAP and GPO-PAP; Roche Diagnostics). Serum LDL-C and HDL-C concentrations were determined using commercial homogeneous direct measurement methods (Roche Diagnostics). All assays were performed in an automated clinical chemistry analyzer (Hitachi/ Roche Diagnostics) at the Institute of Laboratory Medicine, University Hospital Leipzig.

All blood samples were taken in the morning after an overnight fast and stored at −80°C until analyses. From the 1040 Sorbs enrolled in the cohort, a subgroup of 376 subjects was genotyped for haptoglobin and provided blood samples for zonulin measurements (Table 1). Main metabolic characteristics of the study subjects are summarized in Table 2.

**Table 1:**
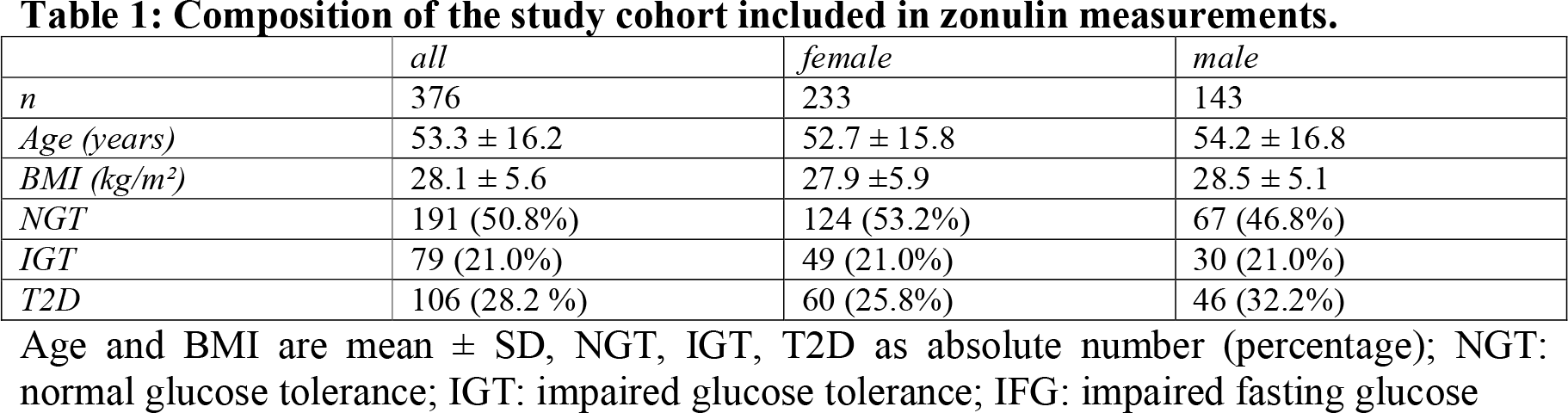
Composition of the study cohort included in zonulin measurements.

|  | *all* | *female* | *male* |
| --- | --- | --- | --- |
| *n* | 376 | 233 | 143 |
| *Age (years)* | 53.3 ± 16.2 | 52.7 ± 15.8 | 54.2 ± 16.8 |
| *BMI (kg/m<sup>2</sup>)* | 28.1 ± 5.6 | 27.9 ± 5.9 | 28.5 ± 5.1 |
| *NGT* | 191 (50.8%) | 124 (53.2%) | 67 (46.8%) |
| *IGT* | 79 (21.0%) | 49 (21.0%) | 30 (21.0%) |
| *T2D* | 106 (28.2 %) | 60 (25.8%) | 46 (32.2%) |

Age and BMI are mean ± SD, NGT, IGT, T2D as absolute number (percentage); NGT: normal glucose tolerance; IGT: impaired glucose tolerance; IFG: impaired fasting glucose

**Table 2:** Main characteristics of the study participants.

|  | *NGT* | *IGT* | *T2D* |
| --- | --- | --- | --- |
| *n* | 191 | 79 | 106 |
| *Age (years)* | 45.4 ± 15.6 | 59.7 ± 29.4** | 62.9 ± 11.2** |
| *BMI (kg/m<sup>2</sup>)* | 25.8 ± 5 | 29.7 ± 4.6** | 30.97 ± 5.53** |
| *WHR* | 0.84 ± 0.10 | 0.92 ± 0.09 | 0.94 ± 0.09** |
| *FPG (mmol/l)* | 5.15 ± 0.50 | 5.68 ± 0.56** | 7.72 ± 2.43** °° |
| *120 min PG (mmol/l)* | 5.00 ± 1.40 | 8.95 ± 0.98** | 5.78 ± 6.86 °° |
| *Triglycerides (mmol/l)* | 1.29 ± 0.94 | 1.56 ± 0.92* | 1.87 ± 1.24** |
| *Fasting Insulin (pmol/l)* | 35.90 ± 25.98 | 53.56 ± 25.98** | 62.38 ± 43.07** |
| *120 min Insulin (pmol/l)* | 148.39 ± 133.58 | 409.28 ± 232.81** | 182.22 ± 272.34 °° |
| *HDL (mmol/l)* | 1.69 ± 0.40 | 1.56 ± 0.36* | 1.43 ± 0.39** ° |
| *LDL (mmol/l)* | 3.20 ± 0.96 | 3.70 ± 1.06** | 3.33 ± 0.95 ° |
| *Total cholesterol (mmol/l)* | 5.12 ± 1.03 | 5.57 ± 1.18* | 5.27 ± 0.99 |
| *HOMA IR (pmol/l\*mmol/l)* | 1.34 ± 1.06 | 2.30 ± 1.1** | 3.58 ± 2.87** °° |
| *HOMA IS (ratio)* | 1.07 ± 0.76 | 0.67 ± 0.78** | 0.68 ± 1.63** |
| *CRP (mg/l)* | 2.22 ± 4.20 | 3.19 ± 4.54 | 3.87 ± 4.76** |
| *Zonulin (ng/ml)* | 67.25 ± 25.45 | 71.88 ± 29.36 | 81.78 ± 25.31** °° |

\*: p<0.05 vs NGT; \*\*: p<0.01 vs NGT
°: p<0.05 vs IGT; °°: p<0.01 vs IGT
All data given as mean ± standard deviation, NGT: normal glucose tolerance, IGT: impaired glucose tolerance, T2D: type 2 diabetes, CRP: C-reactive protein, HDL: High density lipoprotein, LDL: Low density lipoprotein, FPG: Fasting plasma glucose, WHR: Waist to hip ratio

The study was approved by the ethics committee of the University of Leipzig and all subjects gave written informed consent before taking part in the study.

### ELISA measurements

Circulating zonulin was measured by a competitive ELISA (Immundiagnostik AG, Bensheim, Germany) in serum samples of 376 individuals according to the manufacturer’s protocol. Inter-assay coefficient of variation was 6.5%. When purified proteins zonulin and properdin and the synthetic peptide AT1001 were tested by the ELISA, they were re-suspended in PBS and diluted in Diluent buffer (IDK kit) to reach a final concentration of 5 μg/ml.

### Genotyping

Haptoglobin genotypes were determined by PCR using a method adapted from Koch et al. (17). Briefly, the following two primer pairs were used: A (5′-GAGGGGAGCTTGCCTTTCCATTG-3′) and B (5′-GAGATTTTTGAGCCCTGGCTGGT-3′), as well as C (5′-CCTGCCTCGTATTAACTGCACC AT-3′) and D (5′-CCGAGTGCTCCACATAGCCATGT-3′). The primer pair A/B generates two bands: a 1,757-bp allele 1 specific band and a 3,481-bp allele 2 specific band. The primer pair C/D produces one allele 2 specific band of 349 bp. The combination of the bands allows a reproducible typing of the two common haptoglobin genotypes HP1 and HP2. In contrast, no band is detectable for the rare HP deletion genotype, present in ~0.1% Caucasians (16).

### Antibody capturing experiment

We aimed to isolate the target protein of the ELISA antibody from serum samples utilizing the immobilized anti-zonulin antibodies on the ELISA plates to perform antibody capturing experiments. Based to the manufacturing information, these polyclonal antibodies were raised against an octapeptide sequenced from the zonulin molecule (25). Equal amounts of undiluted serum samples with highest and lowest concentrations of zonulin, as measured using the same ELISA kit, were transferred to separate wells, incubated and washed according to the manufacturer’s protocol. Afterwards we eluted the captured protein(s) by incubation with 50 μl of hot (95°C) SDS sample buffer with β-mercaptoethanol for 5 minutes. The captured protein of high or low zonulin serum samples were pooled (N=8 for high (2 pooled groups) and low (1 pooled group); measured protein concentrations using the zonulin ELISA of the serum samples is given in Supplementary Table 1). Twenty μl of these elution samples (high or low zonulin) were separated by SDS-PAGE using precast Bolt 4-12% Bis-Tris Plus gels (ThermoFisher, Waltham, MA, USA). Proteins were stained using the Pierce silver stain for mass spectrometry (ThermoFisher) or detected by Western Blot.

### Mass spectrometry

To identify proteins isolated from serum samples as described above, bands were excised from silver-stained gels and in-gel digested with trypsin following a standard protocol (26). After enzymatic digestion, the peptide mixtures were immediately analyzed by LC/MS/MS on an U3000 RSLC nano-HPLC system (Thermo Fisher Scientific) coupled to an Orbitrap Fusion Tribrid mass spectrometer (Thermo Fisher Scientific). Samples were loaded onto a pre-column (RP-C8, 300 μm * 5 mm, 5 μm, 100 Å, Thermo Fisher Scientific) and washed with water containing 0.1% (v/v) TFA for 15 min, before the peptides were separated on the separation column (RP-C18, 75 μm * 250 mm, 2 μm, 100 Å, Thermo Fisher Scientific) using gradients from 1% to 40% (v/v) B (45 min), 40% to 85% (v/v) B (5 min) followed by 85% B (5 min), with solvent A: 0.1% (v/v) formic acid (FA) in water and solvent B: 0.08% (v/v) FA in acetonitrile. Data were acquired using data-dependent MS/MS mode where each high-resolution full-scan in the orbitrap (m/z 198 to 1,500; R = 120,000) was followed by high-resolution product ion scans in the orbitrap (higher energy collision-induced dissociation (HCD), 27% normalized collision energy, R = 15,000, isolation window 2 Th) within 5 s, starting with the most intense signal in the full-scan mass spectrum.

Data analysis was performed using the Proteome Discoverer 2.1 (Thermo Fisher Scientific). MS/MS data of precursor ions (m/z range 350-5,000) were searched against the Swissprot Database (version 11/2016, taxonomy human, 20,082 entries) and a contaminant database using Sequest HT. Mass accuracy was set to 5 ppm and 20 mmu for precursor and fragment ions, respectively. Carbamidomethylation of cysteines was set as fixed modification; oxidation of methionines and N-terminal acetylation were set as variable modifications, two missed cleavages of trypsin were allowed. Results were filtered for non-contaminant proteins identified by at least three unique highly confident peptides (peptide FDR ≤ 1%).

### Western blot analysis

Western blot experiments were performed to validate the results of mass spectrometric data analysis and to compare the serum target proteins identified by the polyclonal antibodies used by this ELISA kit compared to monoclonal anti-zonulin antibodies. Gels were blotted on a PVDF membrane and Western blots were probed with anti-C3 β-chain (1:2000) (Biozol, Eching, Germany), anti-haptoglobin (1:1000) (Abcam, Cambridge, UK), polyclonal anti-zonulin (1:500) (kindly provided by Immundiagnostik), monoclonal anti-zonulin (1:5000) (BioRad, Hercules, CA USA) antibodies. Purified C3c from plasma (Athens Research, Athens, Georgia, USA) and recombinant zonulin were used as positive controls. Properdin (R&D Systems Minneapolis, MN, U.S.A.) was also used to validate the potential protein candidate identified by the ELISA. Incubation with primary and secondary antibodies (HRP-conjugated) was done for 90 minutes at room temperature. Blots were visualized by enhanced chemiluminescence using Pierce ECL Western Blotting Substrate (ThermoFisher).

### Statistical analysis

Statistical analysis was performed using with SPSS 24 (IBM). All non-normally distributed metric parameters were log transformed to generate a Gaussian normal distribution. Spearman’s rank correlation method was used to assess the relationship between metabolic traits. To test for significant differences in distribution for ordinal values, the Kruskal-Wallis test was used. Exact differences between two groups were tested by both the Mann-Whitney-U test and unpaired student’s t-test. In addition, multiple linear regression analyses were done to assess the linear relationship between continuous variables and genotypes. For all tests, a p-value <0.05 was considered to be statistically significant.

## Results

### Haptoglobin genotype

Haptoglobin genotype HP1/1 was present in 15.8% of the subjects, HP1/2 in 47.6% and HP2/2 in 36.6%. These frequencies are comparable to the distribution of HP genotypes in cohorts of European ancestry reported by others (16, 17). We tested the association of the HP genotypes with various anthropometric and metabolic traits in all study subjects. The analysis revealed that blood hemoglobin levels significantly increase with the presence of at least one HP2 allele (p=0.004 over all three groups, p=4.2×10^−4^ between HP1 homozygote and HP2 carriers). Furthermore, we observed that the total protein concentration in the urine significantly differed between the three groups, with an increase in the HP2 carriers (p=0.027). Interestingly, mean triiodothyronine (fT3) levels were lower in the HP1/1 group than in the HP2/2 group (p=0.012) and in accordance, an increase in administered thyroid gland hormones (p=0.023) was observed.

### Zonulin ELISA data do not match HP genotype distribution

Strikingly, there were no significant differences in levels of the zonulin ELISA signal between the three haptoglobin genotype groups (Figure 1; p=0.153 using ANOVA, p=0.07 for the t-test comparing log transformed zonulin signals between HP1/1 vs. HP1/2 + HP2/2). Assuming that the protein measured by the kit is zonulin (i.e. pre-HP2), subjects with the HP1/1 genotype were expected to have no detectable zonulin levels. As the zonulin concentrations measured in patient sera using the zonulin ELISA kit clearly did not reflect the HP genotype distribution we therefore concluded that the protein measured by the kit is not identical to zonulin as pre-HP2 or that, beside pre-HP2, the kit detects other unrelated proteins. Consequently, we aimed at identifying the protein(s) detected by the alleged zonulin ELISA kit.

**Fig 1.**
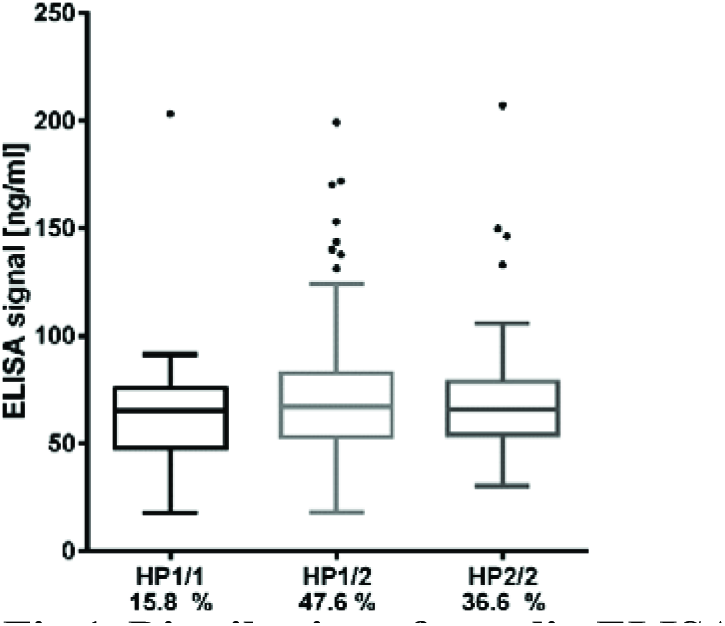
Distribution of zonulin ELISA values according to haptoglobin genotypes. Data is presented as boxplots with Turkey-Whiskers and outliers.

### The zonulin ELISA does not detect recombinant pre-HP2, but targets multiple proteins

To attempt identifying proteins bound and quantified by the capturing antibody deployed in the zonulin ELISA kit, we performed an immune-capturing experiment using patient sera and the immobilized antibody of the ELISA kit as supplied. After incubation of the immobilized ELISA kit antibodies with selected patient sera representing the highest and lowest measured ELISA signals in the cohort, the captured proteins were separated by SDS-PAGE. Notably, we could not measure the protein content of the eluted samples, but given that equal amounts of serum were used, that the same washing and elution procedure was performed for all samples and that 20μl of the pooled elution samples were used for SDS-PAGE and Western blot, the detected amount of “captured” protein should resemble the amount of protein that was present in the initial serum sample.

Silver staining revealed multiple bands, with the most intense band at ~70 kDa and further prominent bands at ~55 kDa, ~180 kDa and >180kDA (Figure 2A). This band pattern was incompatible with a band pattern that would be expected for pre-HP2 or haptoglobin-derived proteins and further supported the results demonstrating the lack of correspondence of the captured protein with HP genotypes. To further characterize major proteins captured by the ELISA kit, protein bands were cut (Figure 2A) and subjected to MS analysis after tryptic digestion (Supplementary Table 2). Mass spectrometry demonstrated that bands 1 (>180kDa), 2 (~150 kDa), 3 (~70 kDa) were all very likely representing the C3 protein or cleavage products derived from the C3 protein, such as the C3 precursor (187 kDa), C3c (144 kDa) and the C3 β-chain (71 kDa). Furthermore, the 55 kDa band was identified as properdin or factor P (MW 53 kDa). To validate results from mass spectrometry, we performed Western blot analyses. The major band at 70 kDa was clearly detected by an anti-C3 β-chain antibody (Figure 2B).

**Fig 2.**
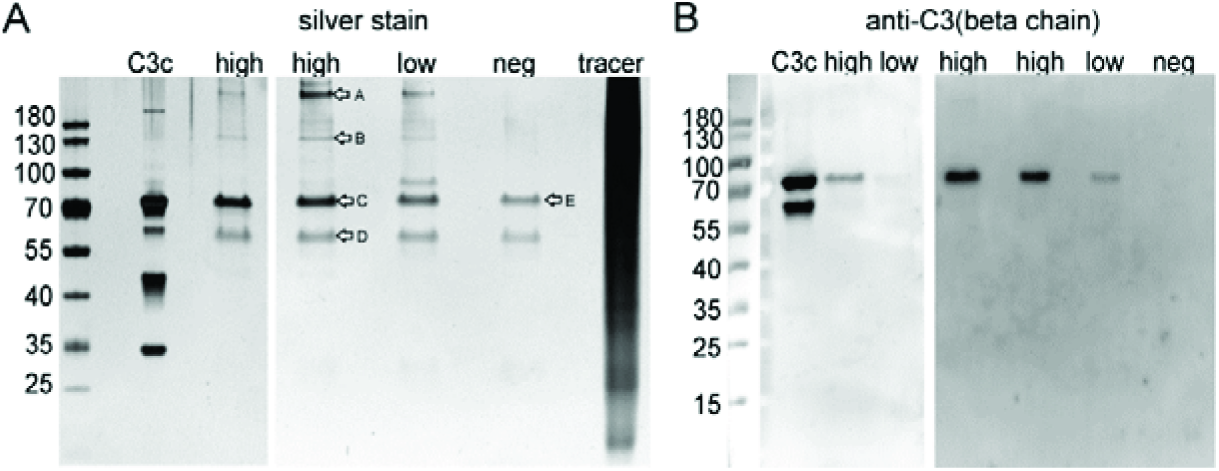
A) Silver stain of immunoprecipitated ELISA products and B) Western blot analyses using an anti-C3 β-chain antibody. Commercially available C3c protein isolated from human plasma was run as positive control. high: pooled IP samples of sera that gave highest ELISA signals; low: pooled IP samples of sera that gave lowest ELISA signals; neg: negative control using dilution buffer from the ELISA kit; tracer: competitive tracer reagent from the ELISA kit. Letters in (A) indicated bands analyzed by MS after tryptic in gel digestion (Supplementary Table 2).

Consequently, we tested several C3c proteins from different suppliers (Abcam, Cambridge, UK; Athens Research, Athens, Georgia, USA; mybiosource, San Diego, California, USA), in a range from 0.1 mg/ml to 0.1 ng/ml under native and denatured conditions, as well as diluted in serum, with the respective ELISA kit. All results were negative (data not shown), indicating that C3 might represent a contaminant only. Additionally, we tested C3, recombinant zonulin, HP1, and HP2 at increasing concentrations (range 1-50 μg/ml), along with sera from celiac patients (both HP1-1 and HP2-2), healthy controls (both HP1-1 and HP2-2), and our standard control (AF, HP2-2). The results showed in Figure 3 demonstrated that while this kit does not recognize C3, zonulin, or mature haptoglobin (both HP1 and HP2), it does recognize protein(s) both in HP1-1 and HP2-2 genotype subjects, irrespective of their disease status.

**Fig 3.**
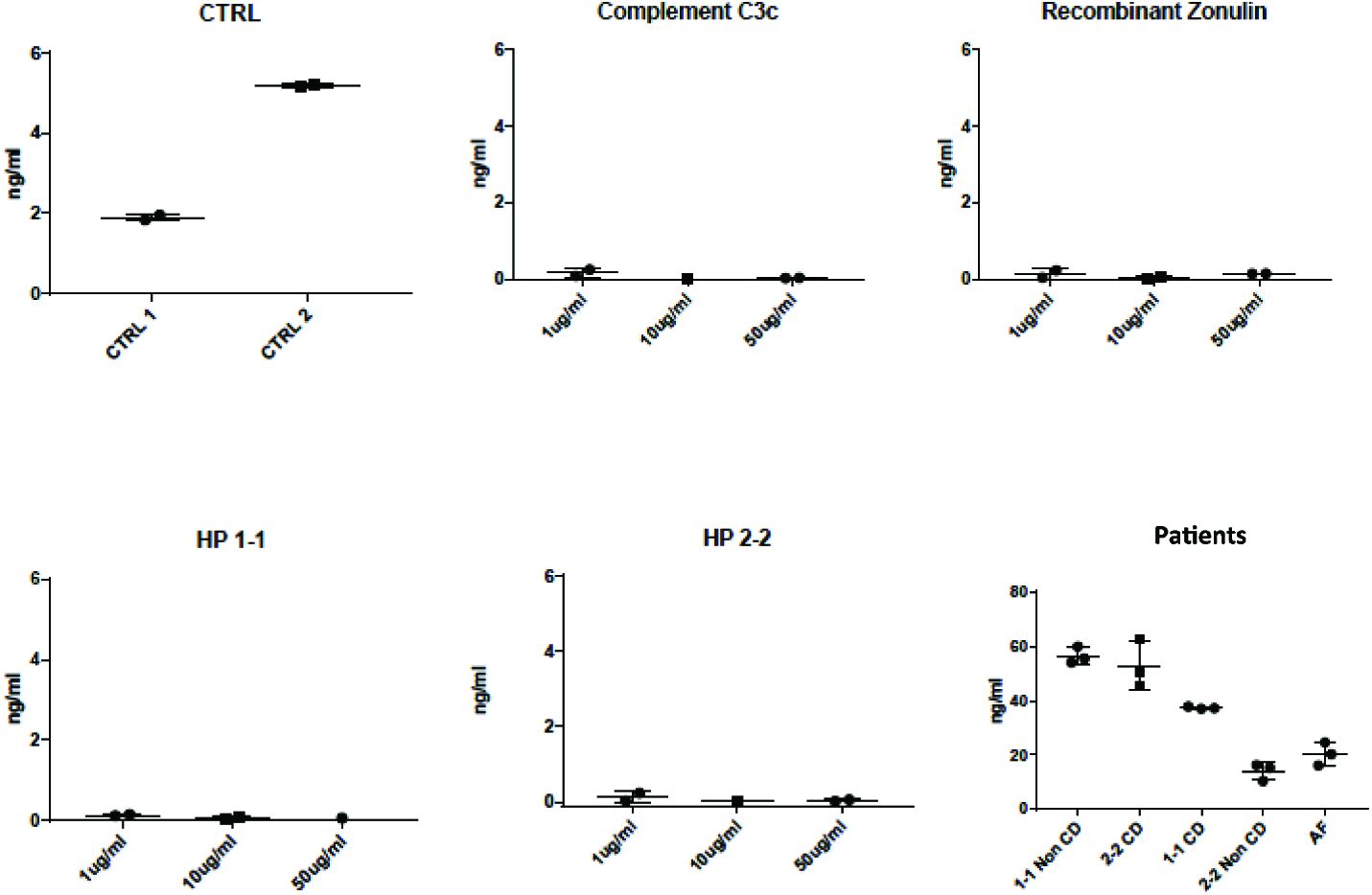
Mean zonulin ELISA values obtained by testing specific candidate target proteins and patients’ serum samples. This ELISA kit did not detect increasing concentrations (range 1-50 μg/ml) of complement, recombinant zonulin (pre-HP2), mature HP1 or mature HP2. Conversely, a strong signal was detected in sera of both celiac disease patients (CD) and healthy controls (non-CD, internal control AF with HP2-2), irrespective of their HP genotype. All samples are technical triplicates, kit controls are shown for comparison.

### Anti-zonulin monoclonal antibodies and Immundiagnostik polyclonal antibodies recognize same target proteins in Western blot

Our ELISA results clearly showed that this kit does not detect zonulin (alias, pre-HP2 as originally claimed by the manufacturer) or complement C3, the protein captured by our experiments described above. In order to further characterize the target protein(s) of this kit, we performed Western blotting analysis of sera from a HP2-2 subject either under baseline condition or after deglycosylation as we have previously described (14), using recombinant zonulin as control. As anticipated, the zonulin monoclonal antibodies recognized recombinant zonulin as well as a variety of bands in the serum sample, including bands with a MW of ~70, 52, 37, 27, and 16 kDa (Figure 4A). Based on similar patterns we detected when zonulin was originally cloned (14), we predicted that the ~72 kDa corresponded to the glycosylated HP β-chain, the 52 kDa zonulin, and the 16 kDa the HP2 α-chain. To confirm this interpretation, we performed deglycosylation experiments showing the shift of the 72 kDa β-chain to a lower MW, while, as anticipated, the zonulin band and the HP2 α-chain remained unchanged (Figure 4A). Interestingly enough, the Immundiagnostik polyclonal antibodies raised against the zonulin synthetic peptide inhibitor GGVLVQPG (AT1001) (25) recognized the same main bands detected by the monoclonal antibodies but with different intensity, being the recombinant zonulin and serum α-chain bands fainter compared to the monoclonal antibody signal, while the serum β-chain and serum zonulin-like signals stronger (Figure 4A). When combined to our ELISA results, these data suggest that, while the zonulin monoclonal antibodies specifically detect in serum samples only zonulin at its predicted molecular weight, most likely the polyclonal antibodies used in this kit are possibly detecting “zonulin-like” protein(s) (as suggested by the much more intense signal of the ~52 kDa band compared to the monoclonal antibodies) with similar molecular weight, structure, and possibly function.

**Fig 4.**
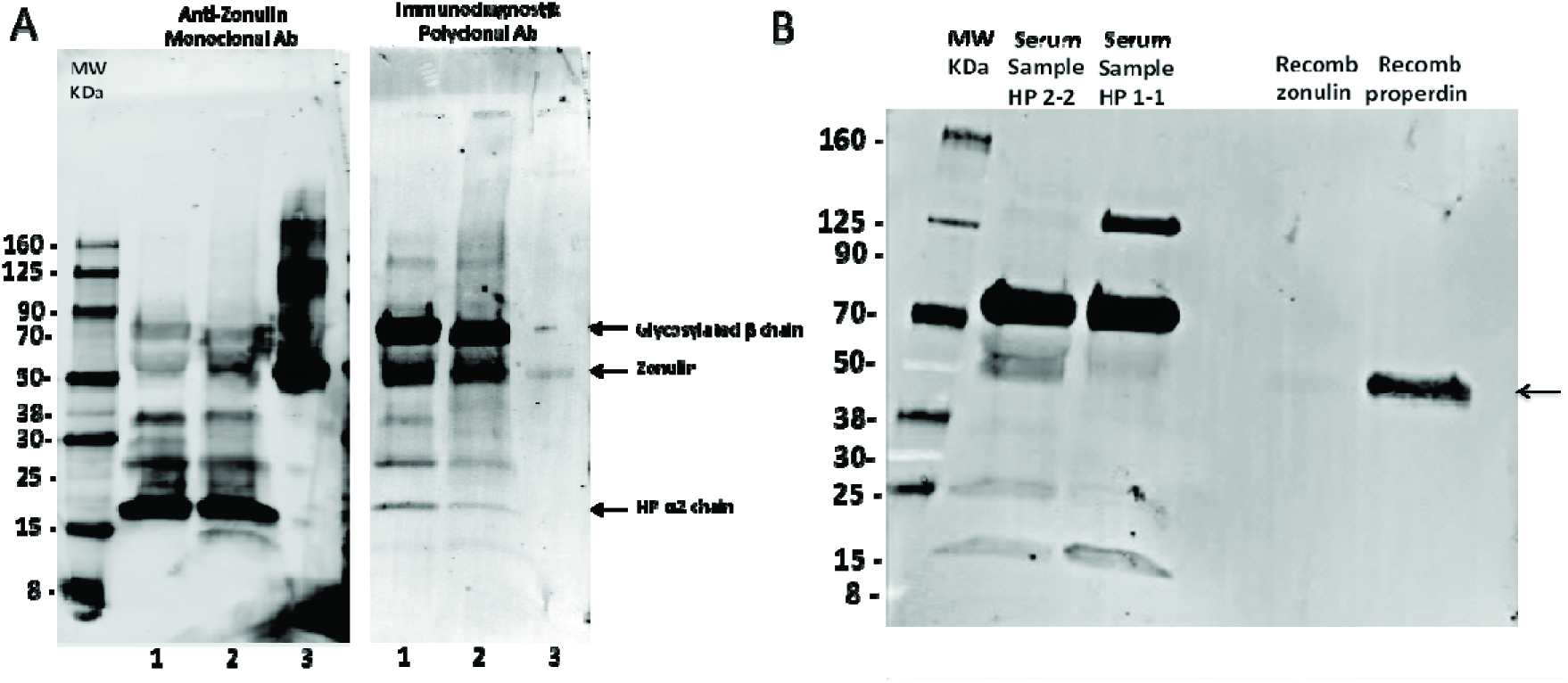
Western blot analyses of a prototype human serum sample (AF HP2-2) using both anti-zonulin monoclonal antibodies and Immundiagnostik polyclonal antibodies. **(A)** Prototype serum sample of a HP2-2 homozygous subject, either untreated (lane 1) or after deglycosylation (lane 2) was resolved and then immunoblotted using either anti-zonulin monoclonal antibodies (left panel) or Immundiagnostik polyclonal antibodies (right panel). Recombinant zonulin was added as control (lane 3). As anticipated, zonulin monoclonal antibodies recognize the recombinant protein as well as a serious of serum bands, with the strongest signal being related to the 16kDa zonulin β-chain. The Immundiagnostik polyclonal antibodies also recognize recombinant zonulin but with a much weaker signal compared to the monoclonal antibodies. These antibodies also recognize the same serum bands detected by the zonulin monoclonal antibodies, with the ~70 and 52 kDa bands being highlighted with the strongest signal. Sample’s deglycosyltation showed the shift of the 70 kDa band to a lower MW, suggesting that this may represent the zonulin β-chain as we have previously shown (14). (B) Prototype serum samples of a HP1-1 and a HP2-2 homozygous subject were resolved and immunoblotted using the Immundiagnostik polyclonal antibodies. Recombinant zonulin and properdin were added as control. The antibodies also detect properdin that migrated at the same molecular weight of zonulin and serum bands recognized by the antibodies in both HP2-2 and HP 1-1 subjects.

### Immundiagnostik polyclonal antibodies recognize both zonulin as well as properdin as an additional target in Western blot

Among all the proteins we identified with our mass spec analysis, properdin fulfills the structural-functional characteristics mentioned above and, therefore, may represent the most likely candidate target detected by this ELISA kit. To explore this hypothesis, we performed both Western blotting analysis using the Immunodiagnostik polyclonal antibodies and ELISA test using commercially available properdin. The Western blot showed that beside recombinant zonulin, the Immunodiagnostik polyclonal antibodies also detect properdin (Fig 4B) that migrated at the same molecular weight of zonulin and serum bands recognized by the antibodies in both HP2-2 and HP 1-1 subjects (Fig 4B). Similar results were obtained with polyclonal antibodies raised against recombinant zonulin (data not shown). The same samples used for Western blotting analysis, including zonulin, properdin and AT1001 at a concentration of 5 μg/ml were tested using the ELISA kit. The results showed that properdin and AT1001 were both detected by the ELISA test, however, their amounts were substantially underestimated by 914 (5.47 ng/ml) and 40 folds (126.04 ng/ml), respectively. However, despite zonulin being detected in Western blot, it was not detected with the ELISA kit (Figure 3). In comparison, serum from subjects with HP 1-1 and HP 2-2 genotype were measured at a concentration of 67.54 and 56.41 ng/ml, respectively (Table 3).

**Table 3:** Correlation of ELISA signal with metabolic phenotypes.

|  | *Non-adjusted* |  | *Adjusted for age, sex, BMI* |  |
| --- | --- | --- | --- | --- |
|  | *r* | *p-value* | *r* | *p-value* |
| *Anthropometric trait* |  |  |  |  |
| *Age* | 0.31 | 0.55 | - | - |
| *Sex* | -0.005 | 0.93 | - | - |
| *BMI* | **0.221** | **1.2x10<sup>-5</sup>** | - | - |
| *WHR* | **0.172** | **7.3x10<sup>-4</sup>** | **0.118** | **2.1 x10<sup>-2</sup>** |
| *Glucose metabolism* |  |  |  |  |
| *Glucose (0 min)* | **0.294** | **6.6x10<sup>-9</sup>** | **0.227** | **9.0 x10<sup>-6</sup>** |
| *Glucose (120 min)* | -0.039 | 0.45 | -0.046 | 0.38 |
| *Insulin (0 min)* | **0.173** | **7.7x10<sup>-4</sup>** | 0.079 | 0.13 |
| *Insulin (120 min)* | -0.039 | 0.46 | -0.044 | 0.39 |
| *HOMA IR* | **0.244** | **2.0x10<sup>-6</sup>** | **0.149** | **4.0 x10<sup>-3</sup>** |
| *HOMA IS* | **-0.243** | **2.0x10<sup>-6</sup>** | **-0.145** | **4.9 x10<sup>-3</sup>** |
| *Lipid metabolism* |  |  |  |  |
| *Triglycerides* | **<u>0.370</u>** | **<u>6.5x10<sup>-14</sup></u>** | **<u>0.312</u>** | **<u>4.2 x10<sup>-10</sup></u>** |
| *Total cholesterol* | **<u>0.219</u>** | **<u>1.5x10<sup>-5</sup></u>** | **<u>0.211</u>** | **<u>3.3x10<sup>-5</sup></u>** |
| *LDL* | **0.182** | **3.4x10<sup>-4</sup>** | **0.160** | **1.7 x10<sup>-3</sup>** |
| *HDL* | **-0.136** | **7.7x10<sup>-3</sup>** | **-0.063** | **0.22** |
| *Apolipoprotein B* | **<u>0.247</u>** | **<u>9.9x10<sup>-7</sup></u>** | **<u>0.215</u>** | **<u>2.3x10<sup>-5</sup></u>** |
| *Adipokines* |  |  |  |  |
| *Adiponectin* | **-0.215** | **2.2x10<sup>-6</sup>** | **-0.176** | **5.4x10<sup>-4</sup>** |
| *Progranulin* | **0.151** | **3.0x10<sup>-3</sup>** | **0.129** | **1.1 x10<sup>-2</sup>** |
| *Vaspin* | 0.027 | 0.6 | 0.05 | 0.33 |
| *Chemerin* | **0.103** | **4.3x10<sup>-2</sup>** | 0.065 | 0.21 |
| *FGF21* | **0.165** | **2.1x10<sup>-3</sup>** | **0.152** | **4.7x10<sup>-3</sup>** |
| *Other* |  |  |  |  |
| *C-reactive protein* | **0.232** | **4.0x10<sup>-6</sup>** | **0.166** | **1.1x10<sup>-3</sup>** |
| *Total protein* | **0.124** | **1.5x10<sup>-2</sup>** | **0.134** | **8.6x10<sup>-3</sup>** |
| *Hemoglobin* | **0.201** | **7.4x10<sup>-5</sup>** | **0.143** | **5.1x10<sup>-3</sup>** |
| *Uric acid* | **0.176** | **5.2x10<sup>-4</sup>** | **0.110** | **3.1 x10<sup>-2</sup>** |

**bold:** significant correlations after adjustment, underlined: significant after Bonferroni adjustment for multiple testing ( $p < 5.2 \times 10^{-4}$ ); r: Spearman rank correlation coefficient, p: significance level; BMI: Body Mass Index; HDL: High density lipoprotein, LDL: Low density lipoprotein, WHR: Waist to hip ratio, FGF21: Fibroblast growth factor 21, HOMA-IR/IS: Homeostasis Model Assessment Insulin Resistance/ Insulin Sensitivity.

### Correlations of measured protein concentrations using the ELISA with metabolic traits

In a sample of 376 subjects tested using the purchased zonulin ELISA, the product was measured in a mean concentration of 72.2 ± 27.2 ng/ml (mean ± standard deviation), ranging from 17.8 to 207.1 ng/ml. The ELISA signal was significantly increased in subjects with T2D (81.78 ± 25.31 ng/ml) compared to subjects with normal glucose tolerance (67.25 ± 25.45 ng/ml, Mann-Whitney-U test; p=2.1×10^−8^) or impaired glucose tolerance (71.88 ± 29.36 ng/ml, p=0.0017) (Figure 5A). Additionally, lean subjects had significantly lower values (65.64 ± 25.23 ng/ml) than subjects with overweight (74.20 ± 30.68 ng/ml, p=0.0082) or obesity (76.24 ± 24.17 ng/ml, p=7.0×10^−5^) (Figure 5B). We observed no gender differences or any correlations with age (data not shown). The ELISA signal correlated with traits related to glucose and lipid metabolism (Spearman’s rank correlation test, adjusted for age, sex and BMI; Table 4). It was positively correlated with BMI, HOMA-IR and -IS and fasting plasma glucose (Table 4). Strong correlations were also observed for lipid metabolism parameters, such as TG levels, total cholesterol, LDL and apolipoprotein B (Table 4). Correlations were tested for a total of 95 accessible traits. After Bonferroni correction for multiple testing (adjusted p-value for significance p < 5.2×10^−4^), correlations for BMI (p=1.2×10^−5^), fasting glucose (p=9.0×10^−6^), TG (p=4.2×10^−10^), total cholesterol (p=3.3×10^−5^) and apolipoprotein B (p=2.3×10^−5^) remained statistically significant.

**Fig 5.**
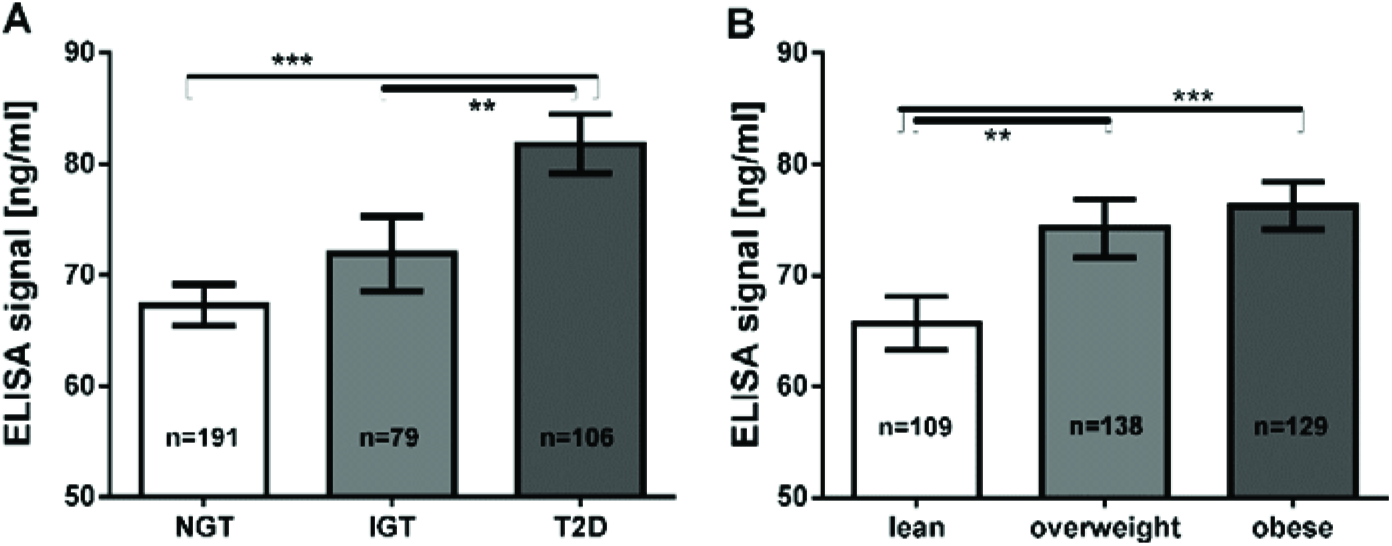
A) Mean zonulin ELISA values according to glucose tolerance groups B) mean ELISA values according to BMI groups. **: p<0.01; ***: p<0.001 NGT: normal glucose tolerance, IGT: impaired glucose tolerance, T2D: type 2 diabetes 696

## Discussion

The role of impaired intestinal permeability in metabolic diseases including obesity and T2D has recently been acknowledged in multiple studies (3, 4, 27). The tight junction regulator zonulin, which was identified as pre-HP2 by Tripathi et al. (14), is an established circulating marker of intestinal permeability in humans (28, 29). Here, we aimed to investigate the relationship between intestinal permeability, represented by circulating zonulin, and metabolic traits related to obesity and T2D. We set out to measure zonulin with a widely used commercially available ELISA kit in a metabolically well-characterized cohort of Sorbs from Germany. Considering the identity of zonulin as pre-HP2, we also genotyped the haptoglobin gene in DNA samples from all subjects. The genotype frequency of the HP1/1 genotype corresponded to previously reported frequencies of 15% in populations of European ancestry (16, 17). Assuming that the measured zonulin is identical with pre-HP2 as reported previously (13), we expected subjects with the HP1/1 genotype to have no detectable, or if taking into account cross-reactions, at least significantly lower zonulin levels. However, concentrations measured in the HP1/1 homozygous subjects were comparable with those found in HP2/2 and HP1/2 carriers.

The epitope used to generate the capture antibody in the zonulin ELISA is based on the sequence GGVLVQPG published by Wang et al. (25) (communication with customer support; Immundiagnostik AG, Bensheim, Germany), which was initially thought to represent the N-terminal sequence of fetal zonulin (25). However, this sequence is not present in pre-HP2, which has since been proposed to be zonulin by the same authors (14). According to the authors, the discrepancies between the previously reported zonulin sequence and the pre-HP2 sequence may be due to intraspecies variability associated with a high zonulin mutation rate or due to sequencing errors at that time (14). In summary, the commercially available competitive ELISA does very likely not detect preHP2 or zonulin, but rather unknown targets bound by the antibody raised against the sequence reported by Wang et al (25). Based on database searches, the epitope could correspond to Glu-Rich2, a protein which shares 7 out of 8 amino acids. The zonulin/preHP2 binding partner CD163 shows some conformity with the epitope (30). Our antibody capture experiment and subsequent mass spectrometry analysis did not provide any evidence for either protein. The most abundant protein identified by MS, C3, is evidently an unspecific product overshadowing the real targets. Indeed, the respective ELISA kit did not detect any complement C proteins obtained from different suppliers when tested under native and denatured conditions, as well as diluted in serum. Also, the same kit did not detect recombinant zonulin, mature HP1 or mature HP2. Considering the additional MS hits, a few proteins stand out, although, without further validation, we interpret these data with caution, since only more abundant proteins may be identified by MS analysis, while our protein(s) of interest may be in low abundance in serum samples and, therefore, not identifiable with this approach. The van Willebrand factor (vWF, band A, Figure 2) is involved in the intrinsic coagulation pathway and the acute phase response and known to be increased in inflammatory bowel disease and bacterial diarrhea (31). Inter-alpha-inhibitor heavy chain 4 (band B, Figure 2) a large glycoportein cleaved into smaller fragments by Kallikrein which is also involved in the intrinsic coagulation pathway. One of these fragments, called urinary trypsin-inhibitor, attenuates LPS-induced endothelial barrier dysfunction by upregulation of vascular endothelial-cadherin expression (32). Complement component 9 (C9, band C, Figure 2) is an important component of the membrane attacking complex within the complement cascade and is required for complement-mediated lipopolysaccharide release and outer membrane damage in bacteria (33). Protein S100-A8, or Calprotectin, (band D, Figure 2) is a calcium -and zinc-binding protein which plays a prominent role in the regulation of inflammatory processes and immune response (34). However, based on the data presented in Figure 4, our most likely candidate(s) should be in the ~50 kDa range, were the polyclonal antibody signal was stronger. Therefore, the most interesting candidate protein we have identified is properdin or factor P (band D, Figure 2), a member of the complement alternative pathway that has a molecular weight (53 kDa), within the range of those proteins recognized by the Immundiagnostik anti-AT1001 polyclonal antibodies (Figures 4A and B) and serum levels (~25 ng/ml) (35) similar to the range of detection of the kit. Our combined Western blot analysis (Figure 4B) and ELISA test confirmed that the polyclonal antibodies raised against AT1001 detect properdin amongst other proteins. However, when purified proteins/peptides, including the AT1001 peptide used to raise the polyclonal antibodies, which is also used as internal control in the ELISA kit, were tested by ELISA, they were highly under-estimated by the test. One possible explanation for these results is that zonulin and also properdin are not the main targets detected by the ELISA, however the fact that even AT1001 was under-estimated seems to suggest that this hypothesis cannot entirely explain our results. Alternatively, it is possible that tertiary and quaternary (multimers) structure arrangements present in sera samples but not in recombinant proteins are necessary in order to properly detect both zonulin and properdin by this ELISA. If this is the case and/or that the main target of this ELISA is/are additional proteins in the ~50kDa range present in human serum remains to be established.

Once released from neutrophils, T cells and macrophages in response to acute microbial exposure, properdin causes production of chemotactic anaphylatoxin C3a and C5a with subsequent formation of immune complexes that cause increased endothelial permeability (35). Intriguingly, zonulin as pre-HP2 also causes generation of C3a and C5a, with subsequent increased vascular permeability in several districts, including the lung, with subsequent onset of acute lung injury (36). Another striking similarity between zonulin and properdin is the fact that both are associated to viral respiratory tract infections (37, 38). Notably, the peptide sequence used for the generation of the antibody is also not present in any of the discussed proteins. The five proteins mentioned above, including properdin, have ~50% similarity to this epitope. Zonulin as preHP-2 is a member of a larger family of tight junction regulating proteins. Indeed, phylogenetic analyses suggest that haptoglobins evolved from mannose-associated serine protease (MASP), a complement-associated protein (like properdin), with their alpha-chain containing a complement control protein (CCP) (this domain activates complement similarly to properdin), while the β-chain is related to chymotrypsin-like serine proteases (SP domain) (39, 40). However, the SP domain of HP lacks the essential catalytic amino acid residues required for protease function; structure-function analyses have implicated this domain in receptor recognition and binding (41). Although not a serine protease, zonulin shares approximately 19% amino acid sequence homology with chymotrypsin, and their genes both map on chromosome 16. Alignment of the β-chain sequence of zonulin to that of several serine proteases is remarkably consistent except for an insertion of 16 residues in the region corresponding to the methionyl loop of the serine proteases. Comparison of the zonulin α-β junction region with the heavy-light-chain junction of tissue-type plasminogen activator strengthens the evolutionary homology of zonulin and serine proteases. The active-site residues typical of the serine proteases, His57 and Ser195, are replaced in zonulin by lysine and alanine, respectively. Because of these mutations, during evolution zonulin most likely lost its protease activity despite that zonulin and serine proteases evolved from a common ancestor (18). Therefore, zonulin, and the serine proteases represent a striking example of homologous proteins with different biological functions but with the common characteristic of complement activation. Beside zonulin and properdin, other members of the MASP family include a series of plasminogen-related growth factors (epidermal growth factor (EGF), hepatocyte growth factor (HGF), etc.) involved in cell growth, proliferation, differentiation and migration, and disruption of intercellular junctions. In light of these considerations, other MASP members identified in our capturing experiments in general, and properdin in particular, are intriguing possible targets (36).

Analyzing the protein concentrations measured using this ELISA in subjects who have been extensively characterized for metabolic phenotypes, our data suggest that it is upregulated both in diabetic and obese patients. This is in line with previously reported findings using this ELISA kit (19–21). Our data shows, that the ELISA target is potentially involved in the lipid metabolism by showing in various linear stepwise regression models that triglyceride levels and fasting glucose are the strongest independent available variables explaining the observed variance in measured protein concentrations (Supplementary Table 3).

It is important to note, that our study, as it has not been initially designed to address the question of ELISA specificity, has clear limitations. To obtain amounts of isolated protein in the antibody capture experiments, we needed to change the experimental protocol provided by the manufacturer. We used undiluted serum samples instead of a 50-fold dilution, which very likely increased the risk of non-specific binding. Yet, we used sera from patient that exhibited the highest and the lowest concentrations measured by the ELISA kit using the manufacturer’s protocol. Thus, any non-specific binding should be detected as equally strong bands in the silver stained gels after the antibody capture experiment. Yet, we observed band intensities of affinity purified protein that clearly correlated with the concentrations measured using the ELISA, in the silver stained gels (total protein) and in the Western blot using the anti-C3-β-chain antibody, indicating a specific isolation of proteins recognized by the kit antibody. Nevertheless, ELISA results that failed to recognize C3 disputed the notion that complement C3 is the target of this kit. Also, we have performed this experiment using two different Lot no. of the ELISA, using sera from eight different patients of each high and low concentration and obtained the same results.

In conclusion, based on our data we suggest that the Immundiagnostik ELISA kit supposedly testing serum zonulin (pre-HP2) levels could identify a variety of proteins structurally and possibly functionally related to zonulin, suggesting the existence of a family of zonulin proteins as previously hypothesized (42), rather than a single member of permeability-regulating proteins. Additional studies are necessary to establish the primary target proteins (zonulin, properdin and/or other structurally similar proteins) detected by this commercially available ELISA.

## Acknowledgements

We thank all those who participated in the studies, in particular our study subjects. We thank Dr. Ingo Bechmann for helpful advice and discussions.

## Author contributions

LS, PK and JTH conceived the study, designed and conducted experiments, analyzed data and wrote the paper. AT recruited patients. CHI performed mass spectrometry experiments. AC, HH, DS and MS interpreted and analyzed data. RF performed the zonulin ELISA test, MF performed the Western blotting analysis, and AF critically revised the manuscript, contributed to the study design of some of the performed experiments, and provided critical interpretation of the data. All authors discussed results, edited and commented on the manuscript. All the authors have accepted responsibility for the entire content of this submitted manuscript and approved submission.

## Supporting Information

**S1 Table: Characteristics of serum samples used for antibody capture experiments**

**S2 Table: Mass spectrometry results of tryptic digests**

**S3 Table: Multiple stepwise linear regression analysis**

